# Time-restricted feeding leads to sex- and organ-specific responses in the murine digestive system

**DOI:** 10.1101/2024.02.20.580958

**Authors:** Lalita Oparija-Rogenmozere, Madeleine R. Di Natale, Billie Hunne, Ada Koo, Mitchell Ringuet, Therese E. Fazio Coles, Linda J. Fothergill, Rachel McQuade, John B. Furness

## Abstract

Food intake is one of the main zeitgebers in the digestive system; however, little is known about organ- and sex-specific differences in food-driven regulation. We placed male and female C57Bl/6 mice on time-restricted feeding (TRF), limiting the food intake period to 8 hours. Food was added either at dark (ZT12) or light (ZT0) onset for 14 days. Afterwards, an additional 4-hour delay in the feeding period was introduced for half of the mice, and the TRF regime continued for another 14 days.

TRF from ZT12 to ZT20 led to the highest weight gain in females but the lowest in males while improving intestinal transepithelial resistance (TEER) in both sexes. However, it also led to the disappearance of food-anticipatory response in several hepatic genes. Delaying the start of TRF until ZT16 led to an increase in weight gain and a decrease in fasting plasma glucose levels in male mice, as well as to strong entrainment of metabolism-related hepatic and duodenal genes in both sexes.

The alignment of food intake with the early lights-on phase (ZT0-ZT8) caused only minor changes in physiological responses. However, it did lead to an overall downregulation of hepatic and an upregulation of duodenal and gastric genes, with additional loss of food-anticipatory gene expression in both sexes. Delaying the start of food intake until ZT4 was highly detrimental, causing an increase in fasting blood glucose levels, a decrease in TEER, and further disruptions in gene expression patterns in the stomach and liver. In contrast, the duodenum was able to restore its food-driven gene expression.

These results demonstrate that the adjustment to food intake time in mice is highly sex- and organ-specific. Our chosen TRF regimes were not able to synchronize food-anticipatory responses between the liver and gut. Instead, we observed that organs entrain to food intake at different rates.

## Introduction

The ability to measure time has played such an importance in our species that clocks are one of the oldest human inventions. We even have devoted an entire subset of science, called Chronometry, to studying timekeeping and developing new methods and standards. And yet, our oldest and arguably most important clock remains understudied.

Self-sustaining and cell-autonomous circadian clocks are essential parts of our biology [1], with early studies revealing diurnal oscillations in the body temperature [2–4], food intake [5], and specific organ functions, for example, renal excretion of water and salts [6, 7]. By now, it has been shown that the circadian clock drives rhythmic expression of approximately 43% of our protein-coding genes and more than a thousand noncoding RNAs [8], regulating major physiological functions across most tissues and organs (for a recent review on the molecular clock and its regulatory role see: [9]).

Simplistically, the mammalian molecular circadian clock is a transcriptional feedback loop, expressed in almost every cell and driving expression patterns of multiple clock-controlled genes with approximately 24-hour rhythmicity. The positive limb of this feedback loop consists of two master transcription factors: CLOCK (circadian locomotor output cycles kaput) and BMAL1 (brain and muscle ARNT-like protein), which form a heterodimer. This heterodimer then binds to the promoters of its target genes, including those comprising the negative limb of the feedback loop: *Per1*, *Per2*, *Per3* (further: PERs – mammalian analogs of the *Drosophila period* gene) and *Cry1*, *Cry2* (further: CRYs). PERs and CRYs dimerize, translocate back to the nucleus, and inhibit the activity of CLOCK-BMAL1 heterodimer, subsequently inhibiting their expression (for extensive reviews on core clock function and mechanism, see [1, 10–14]). There are also multiple secondary loops modulating the output of the core clock, for example, those involving orphan nuclear receptors REV-ERBs and/or retinoic acid receptor-related orphan receptors (RORs). Both of these receptor families have been suggested to provide a link between nutrient metabolism and the circadian clock, especially regarding lipid homeostasis [15, 16]. In addition to the translational regulation, the circadian clock is also regulated post-translationally, with multiple kinases, phosphatases, microRNAs, and RNA-binding protein complexes modulating various aspects and the output of the core clock [1].

Such modulation mechanisms allow for an alignment of circadian rhythms to various cues. In humans, the rhythm can be influenced by age, race, geographical location, and sex [17–20]. In addition, our daily schedules cause recurring variations in environmental and nutritional stimuli, leading to further rhythm adaptation and synchronization. Such stimuli, affecting the timing of any biological rhythm within an organism, are called “zeitgebers” (“time-givers”), a term coined in 1945 [21].

The daily light-dark cycle is the best-researched zeitgeber, entraining the central circadian oscillator that is located in the suprachiasmatic nucleus (SCN) of the hypothalamus. SCN then conveys the generated rhythms to the periphery – directly via neural and hormonal signaling and indirectly via regulation of body temperature. Concurrently, peripheral tissues and organs, for example, the gastrointestinal (further: GI) tract and liver, also possess self-sustaining circadian rhythms with organ-specific phase shifts [22]. These peripheral clocks can entrain to such zeitgebers as food (content and intake time), exercise, and drugs of abuse, potentially leading to their uncoupling from the central clock [23–30]. Circadian misalignment arising from conflicting zeitgeber signals contributes to increased risks of cardiovascular, metabolic, immune, neurological, and even psychiatric disorders (for reviews on health impact, see [31, 32]). In contrast, synchronization between central and peripheral rhythms, for example, by aligning food intake to the light-dark cycle, can be beneficial for general health and well-being (for reviews, see [33, 34]).

Unfortunately, the research on food entrainment of peripheral rhythms is still fragmented. In the liver, food entrainment has been shown to modulate gene expression and metabolic state to account for the anticipated food volume and intake time, even if it requires uncoupling from the SCN-driven rhythm and exacerbating inflammatory responses [23, 35–40]. Ironically, organs comprising the GI tract, which is essential for food uptake, breakdown, and absorption, have been somewhat neglected in food-driven regulation research. Still, several studies have shown that food entrainment regulates GI hormone and short-chain fatty acid secretion [41–43], gut microbiota, lipidome and inflammation [43–45], gastric contractility [43], and expression of clock and clock-controlled genes in small and large intestine [25, 46–49] (for a recent review, see [50]). Synchronization of rhythms across several peripheral organs or tissues in response to food intake time is also poorly studied, with existing reports suggesting that entrainment rates might be organ-specific [23, 46, 51].

Yet another understudied area is sex-specificity in daily rhythms, mostly due to an underrepresentation of female animal models or an overrepresentation of one sex in human studies. There is, however, a physiological basis to expect sex-specific responses, as SCN morphology and neuropeptide expression in rodents differ between males and females [52, 53]. Both rodent and human studies have also shown sex-specific differences in corticosteroid secretion, activity levels, circadian period length, and the stability of the rhythms (for reviews, see [54–56]). When it comes to interaction between sex and food entrainment, male mice seem to develop an earlier and higher-amplitude food-anticipatory behavior when compared to females [57]. In addition, differences in weight gain, clock gene expression, and ghrelin secretion have also been observed [58]. Still, sex-specific differences in food entrainment remain enigmatic.

In this study, we aimed to investigate food-driven, sex-specific regulation of weight gain, glucose handling, intestinal barrier function, and genes involved in nutrient metabolism in the murine digestive system. Adult male and female mice were placed on an 8 h time-restricted feeding (TRF) regime starting either at the onset of the dark (ZT12) or light (ZT0) phase, with or without an additional 4 h feeding delay after 14 days of entrainment. We observed that food intake time not only regulates weight gain, fasting glucose values, and intestinal transepithelial resistance (further: TEER) but also leads to organ- and sex-specific alterations of the core clock (*Clock, Bmal1*) and metabolism-related gene expression.

## Materials and Methods

### Ethical approval

All experimental procedures and handling involving mice were approved by the University of Melbourne Animal Ethics Committee (application #1914983) and performed in compliance with the Australian code for the care and use of animals for scientific purposes.

### Experimental animal origin and housing

In total, 180 male and 180 female C57Bl/6 mice (RRID:IMSR_JAX:000664) from the Animal Resources Centre (Canning Vale, WA, Australia) were used in the study. All mice arrived at the local animal facility aged 5-7 weeks and were kept under a 12:12 h light-dark (further: LD) cycle, with standard chow (Barastoc 102108, Ridley Corporation, Melbourne, Australia) and water provided *ad libitum*. After three days of post-arrival adjustment to the local facility, mice were transferred to either LEDDY cages (black cages with individual lighting systems) (Cat# GM500) or to the Aria ventilated cabinet (Cat# 9BIOC44R4Y1, both from Tecniplast, Buguggiate, Italy), and the light onset of 12:12 h LD cycle was adjusted such that the sample collection from day-fed and night-fed mice could be done in parallel. Mice were adjusted to the new 12:12 h LD cycle and housing conditions for 1 week, based on observations that young mice re-entrain within 5-10 days, especially regarding peripheral rhythms [59] and their activity [60]. Chow and water were provided *ad libitum* during this time. After this week, mice were weighed at the end of the dark phase (Zeitgeber time (ZT) 24). Obtained values were recorded as pre-time-restricted feeding weights (further: pre-TRF) and used as a baseline for later weight gain calculations.

### Food entrainment of mice

After a 1-week adjustment to the new LD cycle, mice were subjected to TRF for 14 days (Fig 1, Pre-shift days). During this time, mice were fed standard chow for 8 h starting either from the dark onset at ZT12 (Fig 1, Restricted night-fed, grey bar) or the light onset at ZT0 (Fig 1, Restricted day-fed, yellow bar), followed by a 16 h fasting period. To ensure that mice did not feed on chow debris during fasting, TRF was done by using two cages: a chow-containing cage (“food cage”) and a cage that never contained any chow (“empty cage”). Mice were placed in the “food cage” during the feeding period and in the “empty cage” during the fasting period. Water was always provided *ad libitum* for the whole duration of the experiment. Mice were weighed weekly at the end of their respective feeding period (Fig 1, blue arrows). Food intake was not measured, as mice were group-housed.

**Fig 1.**
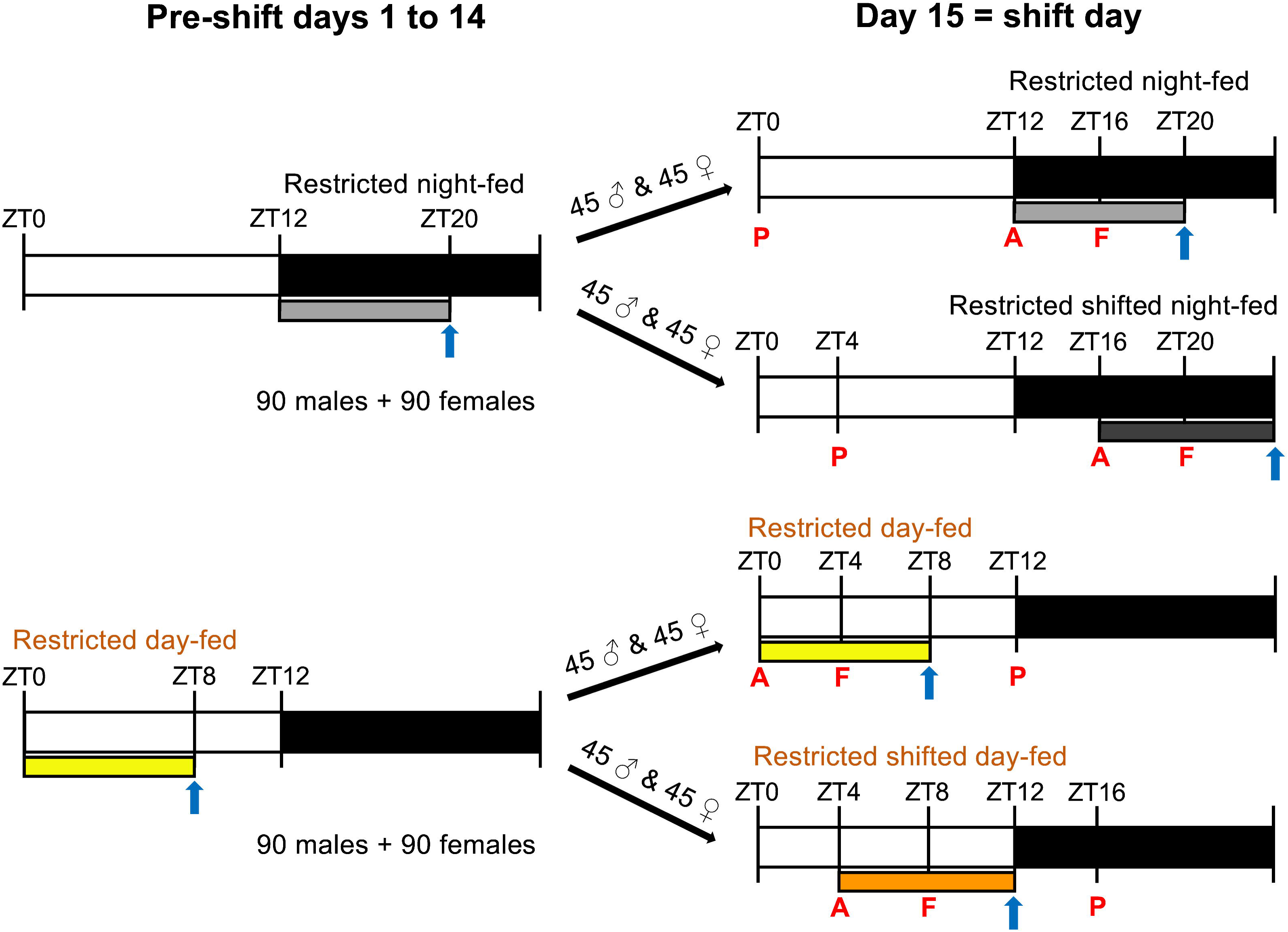
Food entrainment schedule. Bars indicate the placement and length of the feeding period. Arrows show weekly weight measurement timepoints. Mice were sacrificed on the 3^rd^, 7^th,^ or 14^th^ day after feeding shift. Letters indicate sacrifice timepoints on each day: A = food anticipation, F = food intake, P = postprandial. n = 5 per sacrifice timepoint per sex.

After 14 days of the initial entrainment, the 8-hour feeding period was delayed by 4 h for half of the mice (Fig 1, Day 15). The new feeding groups were labeled as ‘Restricted shifted night-fed’ (Fig 1, ZT16-ZT0, dark grey bar) and ‘Restricted shifted day-fed’ (Fig 1, ZT4-ZT12, orange bar). The other half of the mice continued with their initial feeding regimen. Weekly weight gain measurements were continued as previously stated (Fig 1, blue arrows). On the 3^rd^, 7^th,^ and 14^th^ day after the shift, mice were sacrificed for organ collection. Three timepoints were chosen to assess food-driven regulation on each sacrifice day: 1) food anticipation: after 16 h of fasting, right before the anticipated transfer to the “food cage” (Fig 1, “A”); 2) food intake: 4 h after the mice were placed in the “food cage” (Fig 1, “F”); 3) postprandial period: 4 h after the mice were placed in the “empty cage” (Fig 1, “P”). On each sacrifice day, n=5 female and n=5 male mice were sacrificed at each timepoint.

Mice were anesthetized with isoflurane (Cat# FGISO0250, Pharmachem, Eagle Farm, Australia), followed by a cervical dislocation. Upon sacrifice, their body weight was measured, and the following organs and tissues were collected and snap-frozen using liquid nitrogen: the right lobe of the liver, the dorsal half of the stomach, and the duodenal mucosa. To collect duodenal mucosa, the first 2 cm of the duodenum were opened and scraped with a surgical scalpel at a 45-degree angle. The ileum (5 cm proximal from the cecum) was collected only during the food intake timepoint (Fig 1, “F”) and used to measure intestinal permeability and TEER.

### Blood glucose measurements

Blood glucose was measured at the food anticipation timepoint after 16 h fasting (Fig 1, “A”). A drop of blood from the tail vein was applied on the Accu-Chek® Performa Test strip (Cat# 06454038020), and glucose concentration was measured with Accu-Chek® Performa (Cat# 05894964014) blood glucose meter (both from Roche Diagnostics, Manheim, Germany).

### Intestinal electrical resistance measurements

The ileum collected during the food intake timepoint (Fig 1, “F”) was cut in half. Both pieces were placed in Krebs-Henseleit buffer (11.1 mM glucose, 118 mM NaCl, 4.8 mM KCl, 1.0 mM NaH_2_PO_4_, 1.2 mM MgSO_4_, 25 mM NaHCO_3_, 2.5 mM CaCl_2_, pH 7.4), opened along the mesenteric border and pinned (full-thickness) onto Ussing chamber sliders (Cat# P2311, 0.3 cm^2^ apertures, Physiological Instruments, San Diego, USA). Sliders were inserted in the middle of two-part Ussing chambers (EasyMount Diffusion Chambers, Physiologic Instruments, San Diego, USA), and 5 ml Krebs-Henseleit buffer solution was added to the serosal side of the tissue. The mucosal side of the tissue received 5 ml of modified Krebs-Henseleit buffer, where glucose was substituted with 11.1 mM mannitol. This was done to avoid apical uptake of glucose while still maintaining an osmotic balance. Buffers in both chambers were kept at 37°C and bubbled with carbogen (5% CO_2_, 95% O_2_) to maintain an optimal pH level. A multichannel voltage-current clamp (VCC MC6, Physiologic Instruments, San Diego, USA) was applied to each chamber through a set of four electrodes (2 voltage sensing and 2 current passing electrodes) and agar bridges (3% agarose/3 M KCl in the tip and backfilled with 3 M KCl), installed on opposite sides of the tissue. The tissue was left to equilibrate for 20 min before clamping the voltage to 0 V.

To calculate TEER (Ω·cm^2^), 2-sec pulses of 2 mV were administered to tissue every 60 sec for 1 h, and the measured net resistance was multiplied by the surface area. Voltage and short circuit current (Isc) measurements were recorded using a PowerLab amplifier and the software LabChart® 5 (RRID:SCR_017551, both ADInstruments, Sydney, Australia).

### RNA extraction

Total RNA from approx. 20 mg of the right lobe of the liver, dorsal half of the stomach, and scraped duodenal mucosa was extracted using ISOLATE II RNA Mini Kit (Cat# BIO-52073, Meridian Bioscience, Cincinnati, USA) with the following adjustments to the manufacturer’s instructions. First, tissue pieces were immediately snap-frozen in 2 ml screw-cap tubes containing Lysing Matrix D (Cat# 6540-434, Lot# 99999, MP Biomedicals, Irvine, USA) and stored at - 80°C until further processing. Second, the volume of the provided lysis buffer was increased to 500 μl per sample, and 5 μl of β-mercaptoethanol was added. Lysis buffer was applied while the samples were still frozen and immediately followed by lysis at 6000 rpm for 2 x 30 sec in a Precellys 24 homogenizer (RRID:SCR_022979, Cat# 03119-200-RD010, Bertin Technologies SAS, Montigny-le-Bretonneux, France). The volume of RNAse-free Ethanol (Cat# EA043, Chem-supply, Gillman, Australia), used for RNA binding, was also increased to 500 μl per sample, and lysate-ethanol filtration through the provided column was done in two consecutive steps. The centrifugation time for all washing steps was increased to 1 min to ensure better removal of wash buffers. Samples were eluted in 100 μl (liver) or 60 μl (stomach, duodenal mucosa) of RNase-free water (Cat# 10977-015, Lot# 2186758, Invitrogen by Life Technologies, Grand Island, USA). The elution step was repeated using the initial eluate to increase the RNA yield. The quality and quantity of the extracted RNA were assessed using a 2200 Tape Station (RRID:SCR_014994, Agilent Technologies, Santa Clara, USA) and a Nanodrop ND-1000 UV spectrophotometer (RRID:SCR_016517, NanoDrop Technologies, Wilmington, USA), respectively. The isolation was randomized and done in small batches (24 samples) to ensure fast processing and good RNA quality.

The synthesis of cDNA was performed using 100 ng RNA in a 20 μl total reaction volume using an iScript Reverse Transcription Supermix kit (Cat# 1708841, Bio-Rad, Hercules, USA) without any changes from the manufacturer’s instructions. Samples were split across 96-well plates, with 6 negative controls included on each plate, these were randomly chosen RNA samples where no cDNA synthesizing enzyme was added. Synthesis reaction was carried out using PCRExpress (Cat# 630-003, Thermo Hybaid, Franklin, USA) or Bio-Rad T100™ (RRID:SCR_021921, Bio-Rad, Hercules, USA) thermal cyclers.

### Determination of gene expression

After cDNA synthesis, 10 μl of each cDNA sample was delivered to the Translational Research Facility within Monash Health Translational Precinct to determine gene expression levels using the Fluidigm Digital Array Integrated Fluidic Circuits [61] (further: IFCs). First, all cDNA samples underwent quality control, where expression of either *Gapdh* (forward primer: TGACCTCAACTACATGGTCTACA, reverse: CTTCCCATTCTCGGCCTTG) or β*-actin* (forward primer: GGCTGTATTCCCCTCCATCG, reverse: CCAGTTGGTAACAATGCCATGT) was tested as SYBR assays using QuantStudio 6 Flex RealTime PCR reader (RRID:SCR_020239, Thermo Fisher Scientific, Waltham, USA). Afterwards, samples underwent pre-amplification according to the manufacturer’s instructions [62]. In short, all 24 TaqMan assays for genes of interest (Table 1) were first pooled and diluted in Tris-EDTA buffer (pH 8.0) to a final concentration of 180 nM per assay. Then 3.75 μl of gene assay mix was added to 1.25 μl of cDNA sample and pre-amplified for 14 cycles using Veriti™ 96-well Thermal Cycler (RRID:SCR_021097, Cat# 9902, Thermo Fisher Scientific, Waltham, USA). Pre-amplified cDNA samples were further diluted 1:5 with Tris-EDTA buffer (pH 8.0) and loaded on the 192.24 Dynamic array IFC (Cat#100-6266, multiple lots used, Fluidigm, San Francisco, USA) together with gene assays, following Fluidigm® 192.24 Real-Time PCR Workflow Quick Reference PN 100-6170 [62]. The qPCR reaction was performed using the Biomark™ HD system (RRID:SCR_022658, Cat# BMKHD, Fluidigm, San Francisco, USA).

**Table 1.**
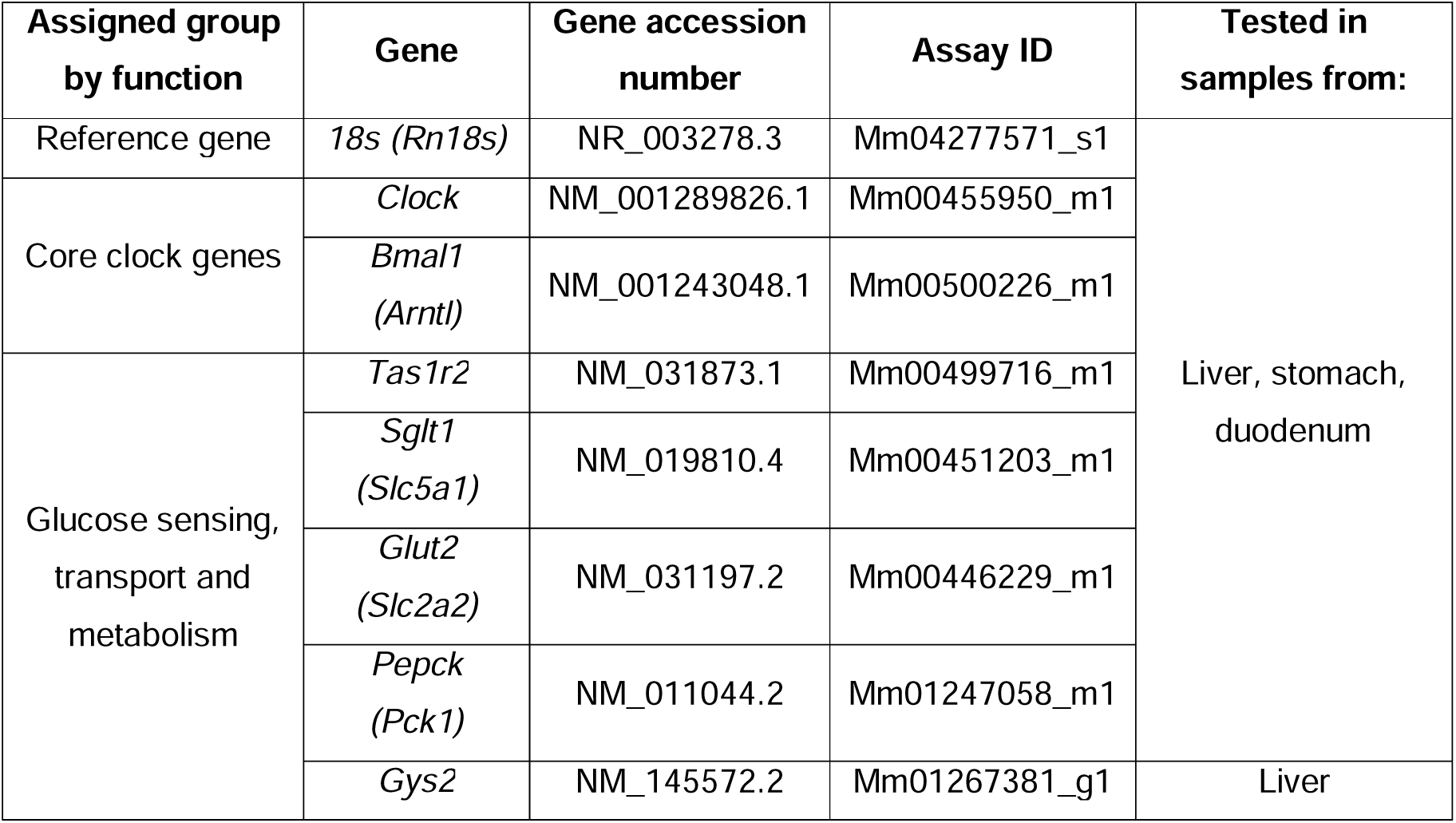
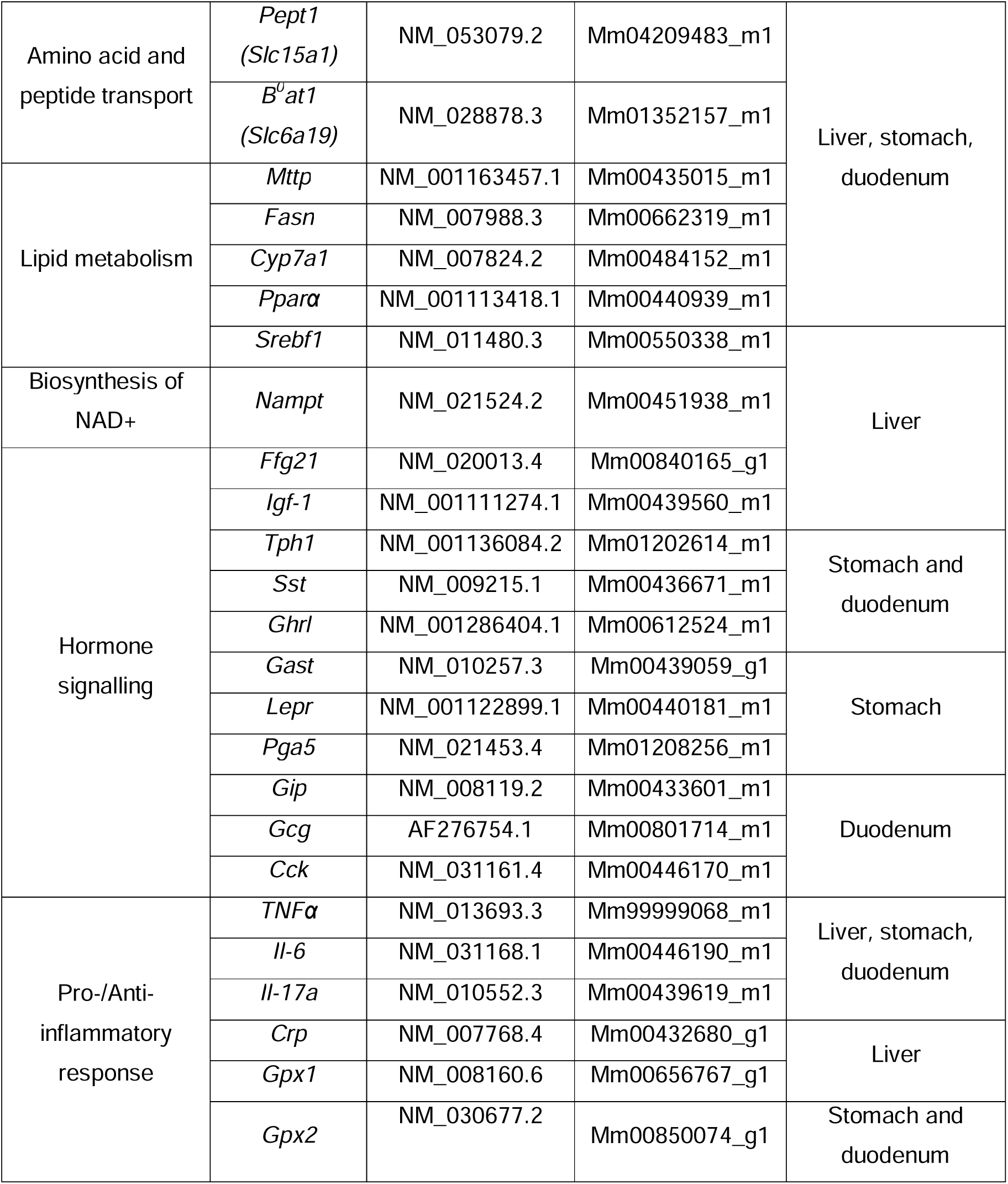
TaqMan Gene assays used to determine relative gene expression levels.

*18s* was used as a reference gene for the calculation of relative expression ratios (R), using the following formula: R = 2^−(Ct(test RNA)-Ct(18S RNA))^. All relative expression ratios were further normalized against the average value shown by night-fed males at the food anticipation timepoint (Fig 1, point “A”) on the 3^rd^ day after the feeding shift.

### Data analysis and statistics

Experimental data analysis was performed using GraphPad Prism v8.2 (RRID:SCR_002798, GraphPad Software, San Diego, CA, USA). To demonstrate weight gain, data from each mouse within the same feeding group and sex (n=90 for pre-TRF, week 1 and week 2; n = 30 for week 3, n = 15 for week 4) were pooled and shown as a mean group value and 95% Confidence Interval, CI. Significant differences in weekly weight gain between the night-fed group and other feeding groups were assessed by Two-way analysis of variance (ANOVA) with Šidák’s multiple comparison test.

Measured TEER values were submitted to an outlier test, with all values outside of mean +/− 2*(standard deviation interval) excluded from further analysis. This was done to exclude tissue pieces that might have been damaged during the resistance measurements.

Weight at the time of sacrifice (Fig 2B), fasting blood glucose values, and TEER values were shown as separate data points per individual mouse, together with the mean group value and standard deviation (Further: SD). Two-way ANOVA was used together with Dunnett’s multiple comparison test to determine differences between all experimental groups independently of the sacrifice day. However, only significant differences between the night-fed group and other TRF regimes on each sacrifice day and the differences between values recorded on various sacrifice days within the same feeding group are shown in the figure.

**Fig 2.**
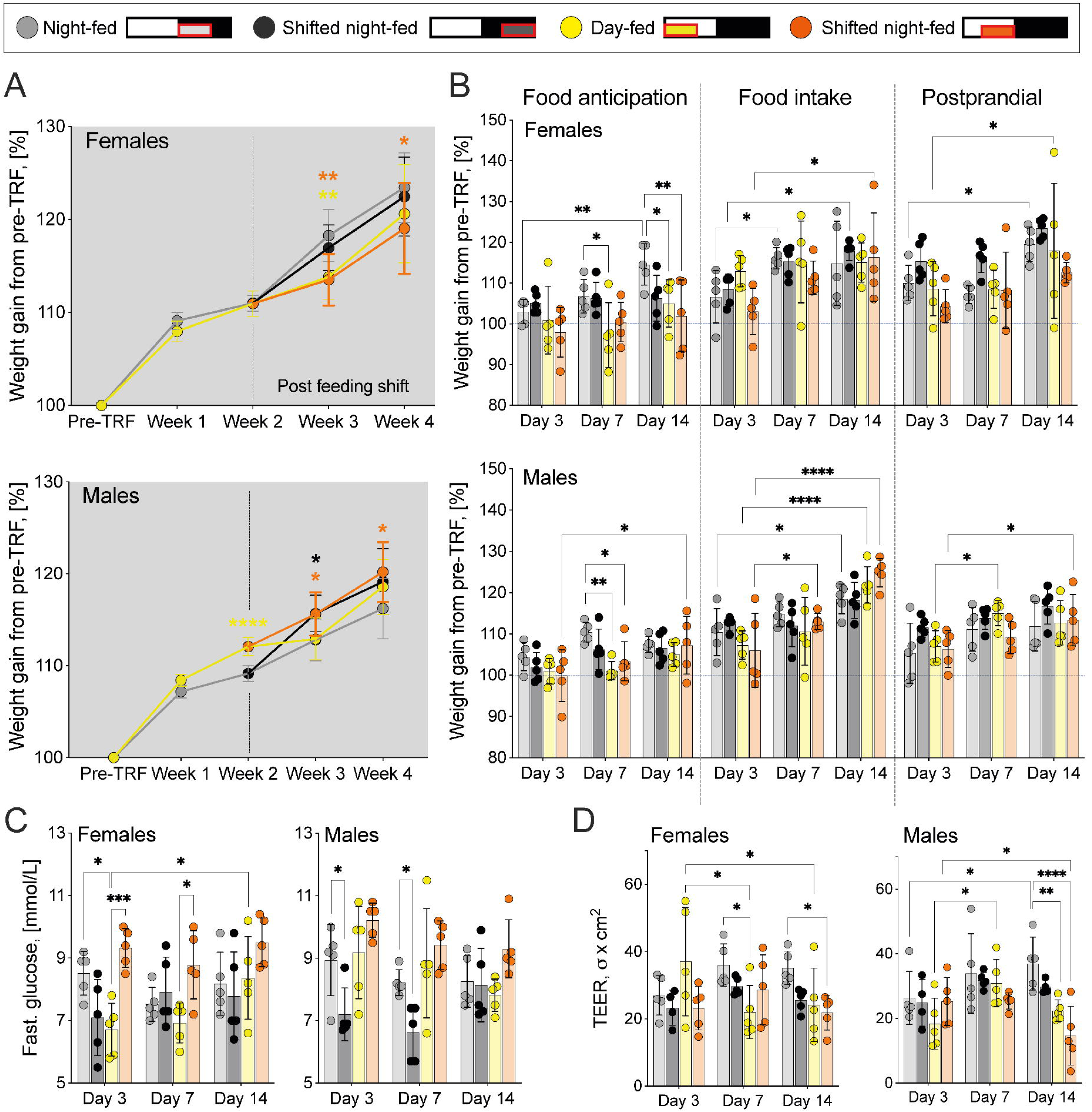
Physiological effects of time-restricted feeding. **A**: Post-feeding weights were normalized to individual pre-food entrainment (pre-TRF) values. Mean (95% CI) shown, n = 90 mice per sex at pre-TRF, week 1 and week 2; week 3 n = 30 mice per sex; week 4 n = 15 mice per sex. Two-way ANOVA with Šidák’s multiple comparison test, showing differences between the night-fed group and other TRF regimes on each sacrifice week, *p<0.05, **p<0.01, ***p<0.001, ****p<0.0001. **B**: Differences in recorded sacrifice weights. Values normalized to individual pre-TRF values. **C**: Fasting glucose values recorded at food anticipation timepoint. **D**: Transepithelial electrical resistance (TEER) in the ileum, recorded at food intake timepoint. **B to D:** Mean (SD) shown, n = 5 mice per sex per timepoint. Two-way ANOVA with Dunnett’s multiple comparison test, showing differences between the night-fed group and other TRF regimes on each sacrifice day and differences between various sacrifice days (Day 3 as a control) within each feeding group, *p<0.05, **p<0.01, ***p<0.001, ****p<0.0001.

To show gene expression differences between male and female mice, normalized expression ratios of mice representing the same feeding group, sacrifice timepoint, and sex were pooled together and shown as a tile within a heatmap. Fold changes in gene expression were shown as Log2 values. To further demonstrate the differences in gene expression between the food anticipation, intake, and postprandial period (=timepoint-specific differences), the nonparametric Kruskal-Wallis test was used together with Dunn’s multiple comparison test (food anticipation timepoint vs food intake or vs postprandial period). The p-values were shown as: *p ≤ 0.05, **p ≤ 0.01, ***p ≤ 0.001, ***p ≤ 0.0001. Differences in gene expression introduced by the 4 h feeding shift were assessed with the same statistical test, comparing values between shifted and non-shifted groups at the same timepoint (e.g., night-fed postprandial vs shifted night-fed postprandial). The p-values were shown as: ^£^p ≤ 0.05, ^££^p ≤ 0.01, ^£££^p ≤ 0.001, ^££££^p ≤ 0.0001.

## Results

### Food-driven regulation of weight gain, fasting plasma glucose levels, and intestinal permeability

#### TRF-induced changes in weight gain are sex-specific

First, all mice had their baseline (pre-TRF) weights recorded. One day later, 180 male and 180 female mice started time restricted-feeding (further: TRF) as either day-fed (further: DF) or night-fed (further: NF) and remained on this regime for 2 weeks. 45 females and 45 males from each TRF regime then underwent an additional 4 h delay shift in the mealtime, creating shifted night-fed (further: SNF) and shifted day-fed (further: SDF) groups. The remaining mice stayed on their initial TRF regime (Fig 1). The total entrainment length was 4 weeks. Weight gain was assessed weekly at the end of the respective feeding period (Fig 1, blue arrows). Afterwards, experimental weights were related to their pre-TRF values and expressed as a change in percentage (%) over time. Weight gain in the NF group was used as a control value in all statistical tests, allowing to determine the impact of lights-on feeding and delay shifts; moreover, this regimen is also the closest to the natural food intake pattern in mice [63].

We observed TRF regime- and sex-specific differences in weight gain. At the end of the 2^nd^ entrainment week, DF males showed a significant 2.9% increase in weight when compared to their NF counterparts. However, this difference disappeared after longer entrainment (Fig 2A, bottom panel, yellow vs grey). In contrast, DF female mice showed significantly delayed weight gain (−4.4%) on week 3 but then also caught up to the NF group (Fig 2A, top panel, yellow vs grey) after longer entrainment.

The additional delay of mealtime caused sex-specific effects as well. In females, we observed similar weight gain between shifted and non-shifted counterparts, but male mice seemed to be more responsive, demonstrating a significantly increased weight gain in both shifted groups one week after the mealtime delay (Fig 2A, in orange vs grey). Overall, it appears that food intake in the late lights-on phase (SNF) was the only condition leading to a similar weight gain pattern in both sexes.

#### Feeding during the lights-on phase leads to weight instability in female mice

The bodyweight was also recorded at each sacrifice timepoint: food anticipation (after 16 h fast, when mice expect to be fed, Fig 1, “A”), food intake (4 h after food was added, Fig 1, “F”) and postprandial period (4 h after food was removed, Fig 1, “P”) on all sacrifice days. The earliest sacrifice was performed on Day 3 after the delayed feeding shift (Day 3), followed by sacrifices on Day 7 and Day 14.

We then compared the sacrifice weights to pre-TRF values (Fig 2B, blue dashed line) and observed that DF and SDF mice showed a more severe weight loss at the food anticipation timepoint, with some even reaching or dipping below the pre-TRF weights. This effect was less pronounced in male mice, where it disappeared under longer entrainment, while females did not show any stabilization. On the contrary – by the end of the experiment, DF and SDF females showed significantly lower food anticipation weights when compared to their NF counterparts (Fig 2B, top panel, Food anticipation).

Weights at the food intake timepoint (Fig 2B, Food intake) tended to increase over time, especially in male mice. There were no differences in weights between various TRF regimes; individual food intake was not measured.

Weights during the postprandial period (Fig 2B, Postprandial) did not differ between TRF regimes, suggesting similar rates of weight loss after 4h fasting.

#### TRF leads to a sex-specific regulation of fasting blood glucose levels

Fasting blood glucose was measured at the food anticipation timepoint (Fig 1, “A”). We observed that female DF mice showed significantly lower fasting blood compared to their NF counterparts (Fig 2C, grey vs yellow). Concurrently, SNF led to significantly lower glucose levels in male mice (Fig 2C, grey vs black). Both of these differences disappeared under longer entrainment and were not present in the opposite sex, suggesting that male and female glucose homeostasis entrains to SNF and DF at different rates. Interestingly, delaying the food intake until the late lights-on phase quickly restored the blood glucose levels in female mice to NF-like-values (Fig 2C, grey vs orange vs yellow), while males showed no difference between DF and SDF.

#### Prolonged food intake during the lights-on phase negatively impacts intestinal barrier function

Ileum from mice sacrificed during the food intake timepoint (Fig 1, “F”) was used to determine TEER. All feeding groups, independently of sex, showed similar TEER values on sacrifice Day 3 (Fig 2D, Day 3). However, we did observe changes under prolonged TRF, especially in male mice. NF males showed an increased TEER over time, while DF and SDF led to a significant decrease by Day 14 (Fig 2D). Female mice showed higher variability, with DF causing significantly lower TEER only on Day 7 but SDF – on Day 14 (Fig 2D). Overall, it appeared that prolonged SDF is detrimental for ileal TEER in both sexes, but NF allows to maintain or even improve the barrier function.

### TRF-driven gene expression in the digestive system

Expression of genes involved in the molecular clock, nutrient metabolism, and immune response was determined in the liver, stomach, and duodenal mucosa. Tissue samples were collected from all experimental animals on sacrifice days 3, 7, and 14 (post-shift), at food anticipation, food intake, and postprandial timepoints (Fig 1, “A”, “F” and “P”, respectively). Gene expression was first normalized against housekeeping gene *18s* and then to the expression level in NF male mice at food anticipation timepoint on sacrifice Day 3. Afterwards, the data from each experimental group (same sex, sacrifice day, and timepoint) was pooled and shown as a single tile within the figure heatmaps. Significant changes in expression between food anticipation and one of the other timepoints within the same TRF group were classified as “timepoint-specific” and shown as *p. Concurrently, significant gene expression differences between shifted and non-shifted groups at the same timepoint were classified as “shift-specific” and shown as ^$^p.

#### Food-driven regulation of *Bmal1* and *Clock* is organ-specific

To determine whether the chosen TRF regimes were able to entrain peripheral core clock genes, we looked at the expression of *Bmal1* and *Clock* in the liver, stomach, and duodenum. We observed that only *Bmal1*, but not *Clock*, showed significant differences between food anticipation and feeding and/or postprandial timepoints in both sexes and all tested organs (Fig 3, *p).

**Fig 3.**
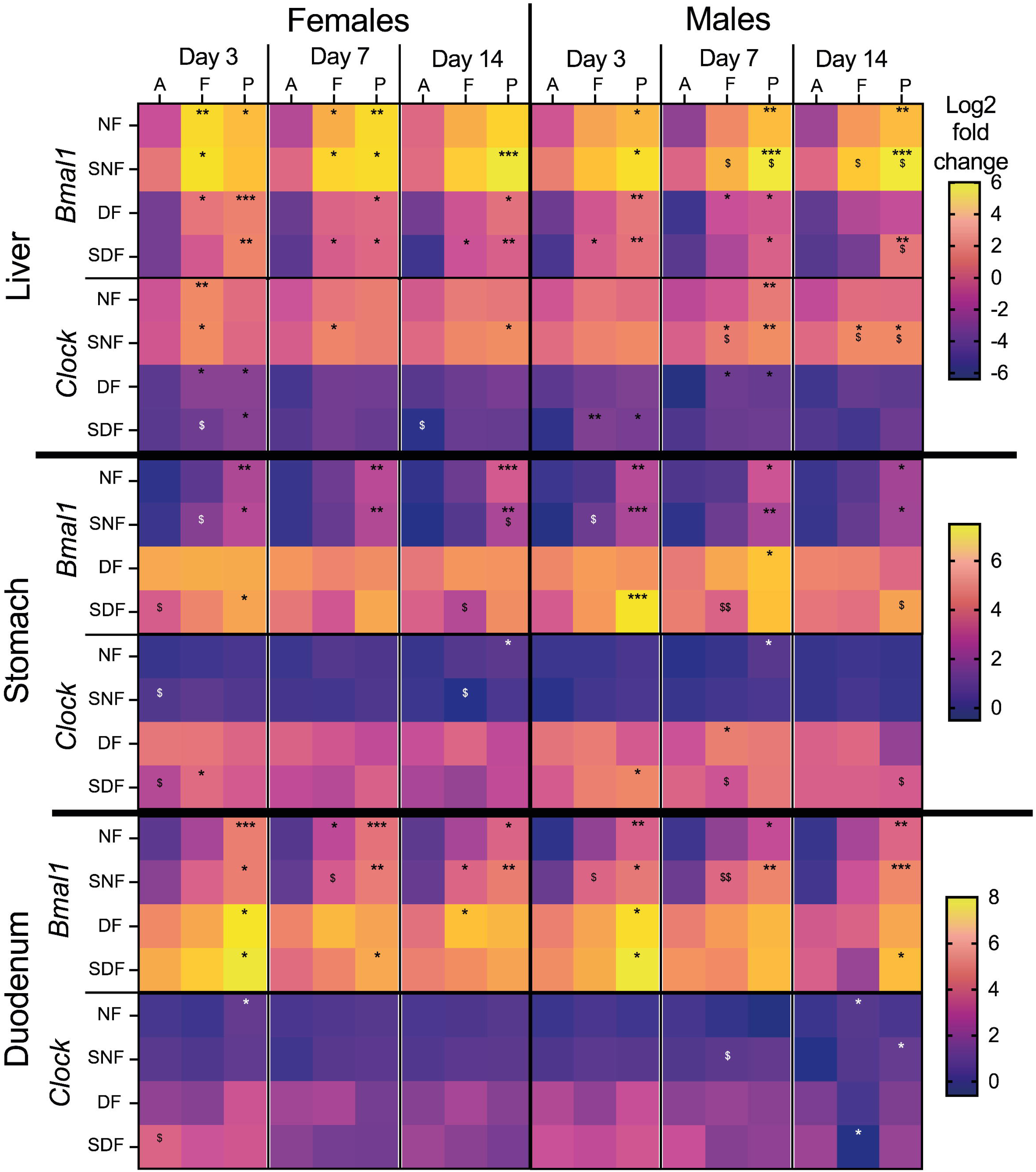
Changes in *Bmal1* and *Clock* expression in male and female mice under TRF. Sample collection timepoints shown at the top: A = food anticipation, F = food intake, P = postprandial period. Feeding conditions shown on the left: NF = night-fed (ZT12 – ZT20), SNF = shifted night-fed (ZT16 – ZT24), DF = day-fed (ZT0 – ZT8), SDF = shifted day-fed (ZT4 – ZT12). Relative gene expression values (Log2) were normalized to NF male mice on sacrifice day 3 at the food anticipation timepoint and pooled by sex, timepoint, and sacrifice day. The mean value of each experimental group is shown as an individual tile (n= 4-5 mice). Non-parametric Kruskal-Wallis test with Dunn’s multiple comparison test used to show timepoint-specific (food anticipation vs intake or vs postprandial period) differences in gene expression on each sacrifice day, *p<0.05, **p<0.01, ***p<0.001. The same test was used to assess shift-specific differences (non-shifted vs shifted) at each timepoint, ^$^p<0.05, ^$$^p<0.01.

It has been shown that in *ad libitum* fed mice under a standard 12:12h light cycle, *Bmal1* expression in the liver reaches its peak in the late dark phase (between ZT20 and ZT0) and the nadir right before the dark onset (between ZT8 and ZT12) [64]. We observed the peak expression of the hepatic *Bmal1* at the food intake or postprandial timepoint under all tested TRF regimes, indicating a response to the food entrainment and possible uncoupling from the light-driven rhythm under DF and SDF conditions. Some timepoint-specific responses were also present for the hepatic *Clock* gene; however, they were not as persistent.

In contrast to the liver, *Bmal1* expression in the stomach and duodenum responded poorly to daytime feeding, with similar expression levels at all timepoints or with the disappearance of changes over time, respectively (Fig 3). However, in NF and SNF male and female mice, we still observed gastric and duodenal *Bmal1* upregulation at the postprandial timepoint. The *Clock* gene did not exhibit recurrent, timepoint-specific changes in expression levels, paralleling previous observations that this gene is not cyclic in the stomach [65].

The 4 h feeding delay had little effect on *Bmal1* or *Clock* expression at tested timepoints, as significant differences between shifted and non-shifted groups appeared rarely and somewhat randomly (Fig 3, differences shown as ^$^p). A more interesting consequence of TRF was the overall lower expression of both clock genes in the liver under DF and SDF regimes (compared to their NF counterparts), while in the stomach and duodenum, we observed the opposite (Fig 3).

Sex-specific differences in gene expression patterns were also present. In some cases, for example, for hepatic *Bmal1* under NF, hepatic *Clock,* and gastric and duodenal *Bmal1* under NF and SNF, it appeared that in female mice, timepoint-specific changes were more likely to be present early (on Day 3 post-shift) and then weaken/disappear over time, while in males the response appeared later, but persisted over longer entrainment.

#### Food intake during the lights-on phase downregulates hepatic gene expression

Similar to our observations with clock genes, also hepatic nutrient metabolism- and inflammatory response-related genes showed an overall lower expression in DF and SDF male and female mice (Fig 4). In addition, even though timepoint-specific expression differences were present for multiple genes, not all of them aligned their expression pattern to every TRF regime tested, leading to an overall poor response to food entrainment.

**Fig 4.**
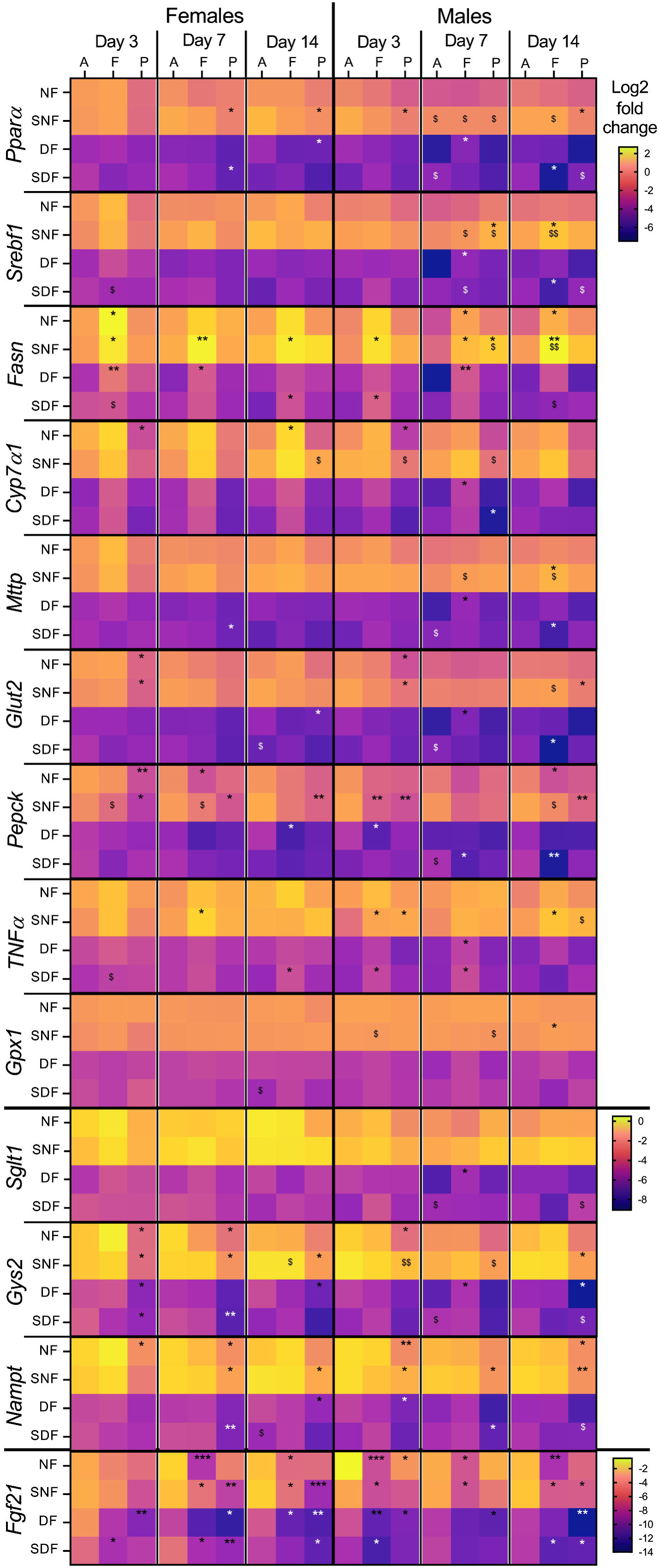
Hepatic gene expression changes in male and female mice under TRF. Sample collection timepoints shown at the top: A = food anticipation, F = food intake, P = postprandial period. Feeding conditions shown on the left: NF = night-fed (ZT12 – ZT20), SNF = shifted night-fed (ZT16 – ZT24), DF = day-fed (ZT0 – ZT8), SDF = shifted day-fed (ZT4 – ZT12). Relative gene expression values (Log2) were normalized to NF male mice on sacrifice day 3 at the food anticipation timepoint and pooled by sex, timepoint, and sacrifice day. The mean value of each experimental group is shown as an individual tile (n= 4-5 mice). Non-parametric Kruskal-Wallis test with Dunn’s multiple comparison test used to show timepoint-specific (food anticipation vs intake or vs postprandial period) differences in gene expression on each sacrifice day, *p<0.05, **p<0.01, ***p<0.001. The same test was used to assess shift-specific differences (non-shifted vs shifted) at each timepoint, ^$^p<0.05, ^$$^p<0.01.

Surprisingly, *Pparα* and *Srebf1*, the principal transcription factors regulating hepatic lipid metabolism [66] that have been observed to be responsive TRF [39, 67], showed discontinuous timepoint-specific changes in gene expression (Fig 4, *p) with only SNF females (*Pparα*) exhibiting downregulation at postprandial period on more than one sacrifice day. The 4h food intake delay also led to only minor changes, mostly by upregulating gene expression in SNF male mice over prolonged entrainment, when compared to their non-shifted counterparts (Fig 4, ^$^p).

However, not every gene associated with nutrient metabolism was so unresponsive to TRF. Genes encoding fatty acid synthase (*Fasn*), glycogen synthase (*Gys2*), and nicotinamide adenine dinucleotide (NAD^+^) biosynthetic enzyme (*Nampt*) showed TRF-induced expression changes, similar to observations before [39]. In addition, responses were also seen from genes encoding phosphoenolpyruvate carboxykinase (*Pepck*) and fibroblast growth factor 21 (*Fgf21*) (Fig 4, *p). Although we observed some sex- and timepoint-specific gene expression differences under DF and SDF regimes for all these genes, the most stable and continuous response in both sexes was seen under SNF. An exception here was *Fgf21*, showing strong downregulation during food intake and/or postprandial period under all TRF regimes, especially under longer entrainment. Since *Fgf21* is involved in the regulation of nutrient and energy homeostasis and has a fasting-induced secretion (for a review on its role, see [68]), it might have been entrained to the daily 16 h fasting period in our mice, resulting in food-anticipatory upregulation. Not much is known about the food-driven regulation of hepatic (anti-)inflammatory markers. Unfortunately, we were not able to clarify it either, as *TNFα* showed an inconsistent upregulation at the food intake timepoint under SNF and SDF, and no timepoint-specific pattern was present in *Gpx1* expression.

Sex-specific differences seemed to follow a pattern observed for clock genes, with females showing early timepoint-specific differences that tended to disappear over time, but males developing late and/or more stable responses (Fig 4., e.g., *Fasn, Pepck, Gys2, Nampt).* Male mice also appeared to be more likely to up- or down-regulate their gene expression in response to mealtime shift, especially after 4 h food intake delay in the dark phase (SNF vs NF, ^$^p, see Fig 4: *Pparα*, *Srebf1, Fasn, Cyp7α1, Mttp, Gys2*).

#### TRF fails to elicit gene responses in the stomach

As a part of the digestive tract, the stomach is directly exposed to the ingested food and has been shown to synchronize its hormone secretion to mealtime (for a review, see [69]). And yet, gastric gene expression in our study was largely unresponsive to TRF (Fig 5), a phenomenon also observed in rats [70]. Only one major response was present in our mice – food intake during the lights-on phase caused an overall gene upregulation (Fig 5, DF, SDF), following the pattern seen for the gastric clock genes (Fig 3).

**Fig 5.**
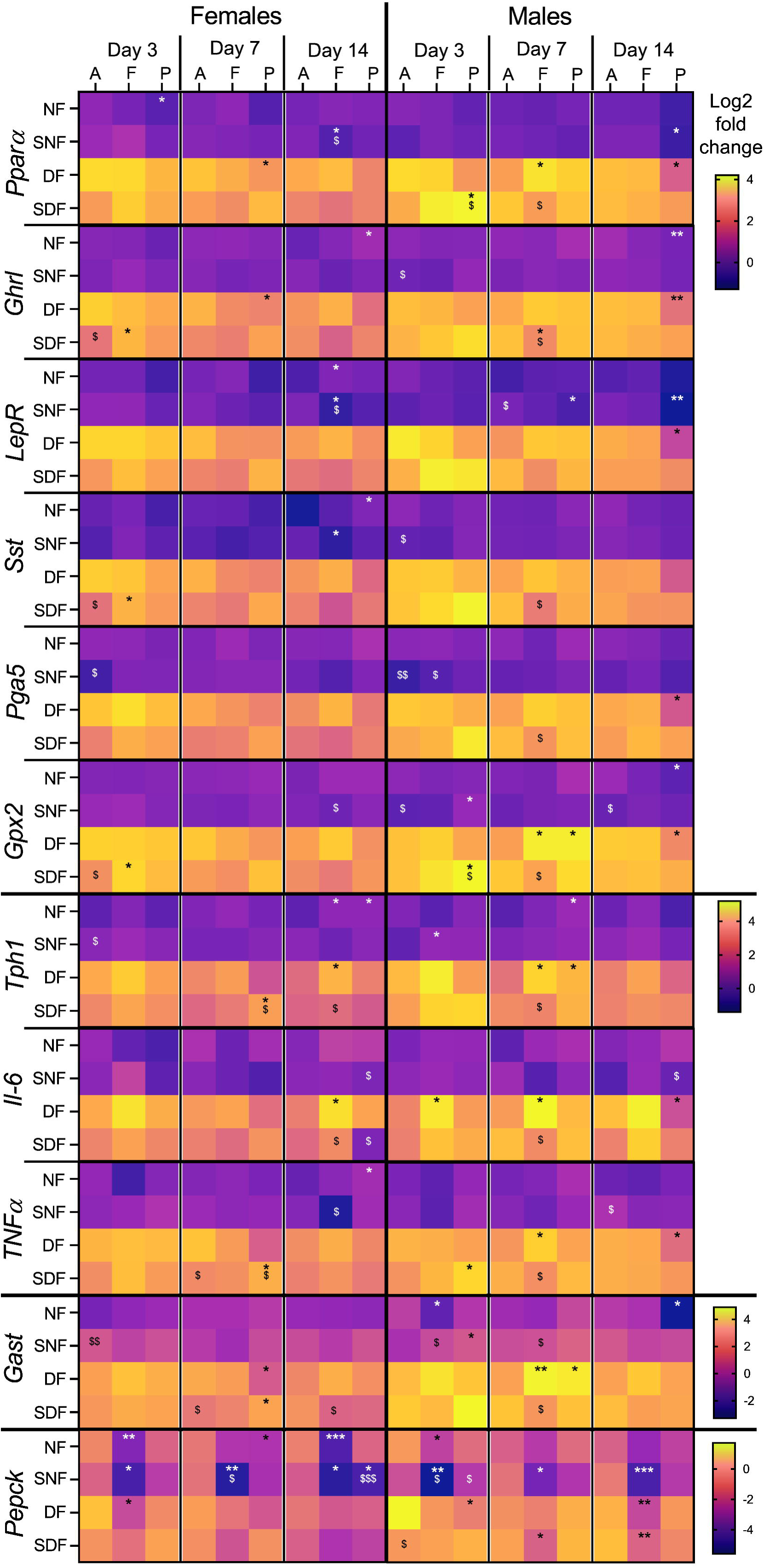
Gene expression changes in the stomach of male and female mice under TRF. Sample collection timepoints shown at the top: A = food anticipation, F = food intake, P = postprandial period. Feeding conditions shown on the left: NF = night-fed (ZT12 – ZT20), SNF = shifted night-fed (ZT16 – ZT24), DF = day-fed (ZT0 – ZT8), SDF = shifted day-fed (ZT4 – ZT12). Relative gene expression values (Log2) were normalized to NF male mice on sacrifice day 3 at the food anticipation timepoint and pooled by sex, timepoint, and sacrifice day. The mean value of each experimental group is shown as an individual tile (n= 4-5 mice). Non-parametric Kruskal-Wallis test with Dunn’s multiple comparison test used to show timepoint-specific (food anticipation vs intake or vs postprandial period) differences in gene expression on each sacrifice day, *p<0.05, **p<0.01, ***p<0.001. The same test was used to assess shift-specific differences (non-shifted vs shifted) at each timepoint, ^$^p<0.05, ^$$^p<0.01, ^$$$^p<0.001.

Despite the lack of stable and continuous TRF-induced changes in expression, multiple genes appeared to follow similar response patterns. For example, genes encoding major GI hormones or their precursors, such as *Ghrl, Pga5,* and *Lepr* (encodes leptin receptor), as well as genes encoding (anti-)inflammatory markers *Gxp2*, *Il-6* and *TNFα* and transcription factor *Pparα*, all showed downregulation at the postprandial period in DF male mice on Day 14 and a considerable overlap in other timepoint-specific changes (Fig 5, *p). A similar expression pattern was seen also for *Tph1,* which encodes tryptophan hydroxylase – a rate-limiting protein in serotonin synthesis.

Gene encoding gastrin (*Gast*), a hormone stimulating gastric acid release and gastric mucosal growth [71], showed a chaotic timepoint-specific gene expression pattern as well, with changes appearing and disappearing between sacrifice days and TRF regimes. Some studies have suggested that *Pparα* could regulate *Gast* expression and/or translation [72, 73]; however, we observed no similarities between their gene responses.

An unexpected observation was the strong timepoint-specific gene expression pattern in gene *Pepck*, which encodes a rate-limiting enzyme for gluconeogenesis. Downregulation during food intake was seen in SNF males and females, as well as in NF females and DF and SDF males after prolonged entrainment (Fig 5). Although gluconeogenesis can occur in the small intestine [74], especially after gastric bypass surgery [75], currently, there is no evidence of glucose production in the stomach, making us wonder why such gene regulation would be necessary. However, *Pepck* has been proposed to play a role also in the regulation of the TCA cycle flux [76] and nutrient processing [77]; therefore, its timepoint-specific regulation in the stomach might be important for gluconeogenesis-independent aspects of nutrient metabolism.

Gene expression changes induced by 4 h mealtime delay were uncommon in the stomach, appearing only on select timepoints and sacrifice days (Fig 5, ^$^p). The lack of stable timepoint- and shift-specific expression patterns was present in both sexes, obscuring any further sex-specific difference detection.

#### Duodenal genes involved in nutrient metabolism respond to food entrainment

The duodenum is responsible for digestion and absorption of nutrients; thus, it would not be surprising if food intake and nutrient-related signals would regulate its gene expression. Indeed, duodenal genes involved in glucose and amino acid uptake and/or metabolism were responsive to TRF (Fig 6). Some similarities to our observations in the stomach were also present: daytime feeding led to an overall gene upregulation, and the genes encoding GI hormones or their precursors (*Cck, Gip, Ccg, Tph1*) showed poor responses to TRF (Figs 5-6).

**Fig 6.**
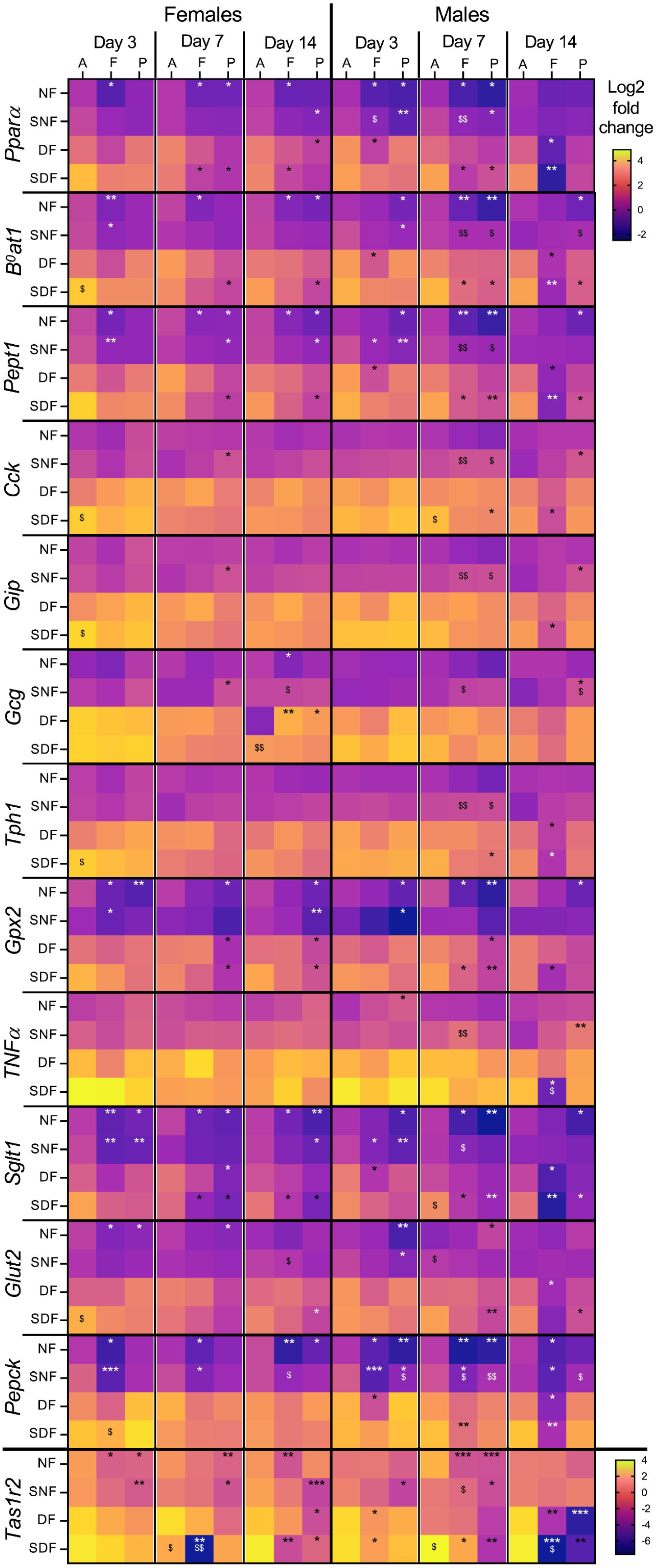
Duodenal gene expression changes in male and female mice under TRF. Sample collection timepoints shown at the top: A = food anticipation, F = food intake, P = postprandial period. Feeding conditions shown on the left: NF = night-fed (ZT12 – ZT20), SNF = shifted night-fed (ZT16 – ZT24), DF = day-fed (ZT0 – ZT8), SDF = shifted day-fed (ZT4 – ZT12). Relative gene expression values (Log2) were normalized to NF male mice on sacrifice day 3 at the food anticipation timepoint and pooled by sex, timepoint, and sacrifice day. The mean value of each experimental group is shown as an individual tile (n= 4-5 mice). Non-parametric Kruskal-Wallis test with Dunn’s multiple comparison test used to show timepoint-specific (food anticipation vs intake or vs postprandial period) differences in gene expression on each sacrifice day, *p<0.05, **p<0.01, ***p<0.001. The same test was used to assess shift-specific differences (non-shifted vs shifted) at each timepoint, ^$^p<0.05, ^$$^p<0.01.

Genes encoding large amino acid (*B^0^at1*), peptide (*Pept1*) and glucose (*Sglt1*) transporters, a sweet taste receptor (*Tas1r2*), an anti-inflammatory marker (*Gxp2*), a gluconeogenesis regulator (*Pepck*) and a transcription factor peroxisome proliferator-activated receptor alpha (*Pparα*) all had similar timepoint-specific expression patterns (Fig 6, *p). Most of these genes were downregulated during food intake and/or postprandial timepoint under NF and SNF in both sexes. Interestingly, while DF did not always lead to timepoint-specific differences (especially in females), SDF was able to induce very similar expression patterns as NF, suggesting successful alignment to food-intake time. It is also interesting that apart from *Bmal1*, *Pepck* was the only other gene responding to TRF in all tested organs and both sexes.

Sex-specific differences were present as well. Females appeared to preserve or even increase the timepoint-specific gene expression changes during prolonged NF, while males showed stronger responses under prolonged DF and SDF. Shift-specific differences seemed to be more prevalent in male mice (Fig 6, ^$^p), especially in the SNF group, peaking on Day 7 post-shift. Some genes (*B^0^at1, Cck, Gip, Tph1,* and *Glut2)* showed upregulation at food anticipation timepoint in SDF females when compared to their non-shifted counterparts; however, this difference was only present on the earliest sacrifice day.

Overall, we observed that different organs entrain to food intake time at various rates. None of the used TRF regimes were able to synchronize timepoint-specific differences across the liver, stomach, and duodenum. In addition, responses were also sex-specific, with male mice appearing more sensitive to 4 h delay shift in the mealtime and female mice often exhibiting earlier but less stable changes in gene expression.

## Discussion

This study describes how food intake time affects weight gain, fasting glucose levels, intestinal permeability, and the expression of genes involved in the molecular clock, nutrient metabolism, and inflammatory processes in the digestive system of male and female mice. We utilized a TRF model to limit the food consumption to 8 h in a 24 h period, with the initial experimental groups starting their feeding period either at ZT12 (NF) or ZT0 (DF). After 14 days of entrainment, food intake time was delay shifted by 4 h for half of the mice (Fig 1), creating two new experimental groups: SNF (ZT16 start) and SDF (ZT4 start).

### TRF impact on weight gain, glucose levels, and intestinal permeability

Observed changes in physiological parameters (relative to the NF group) during the full food entrainment period have been summarized in Table 2.

**Table 2.**
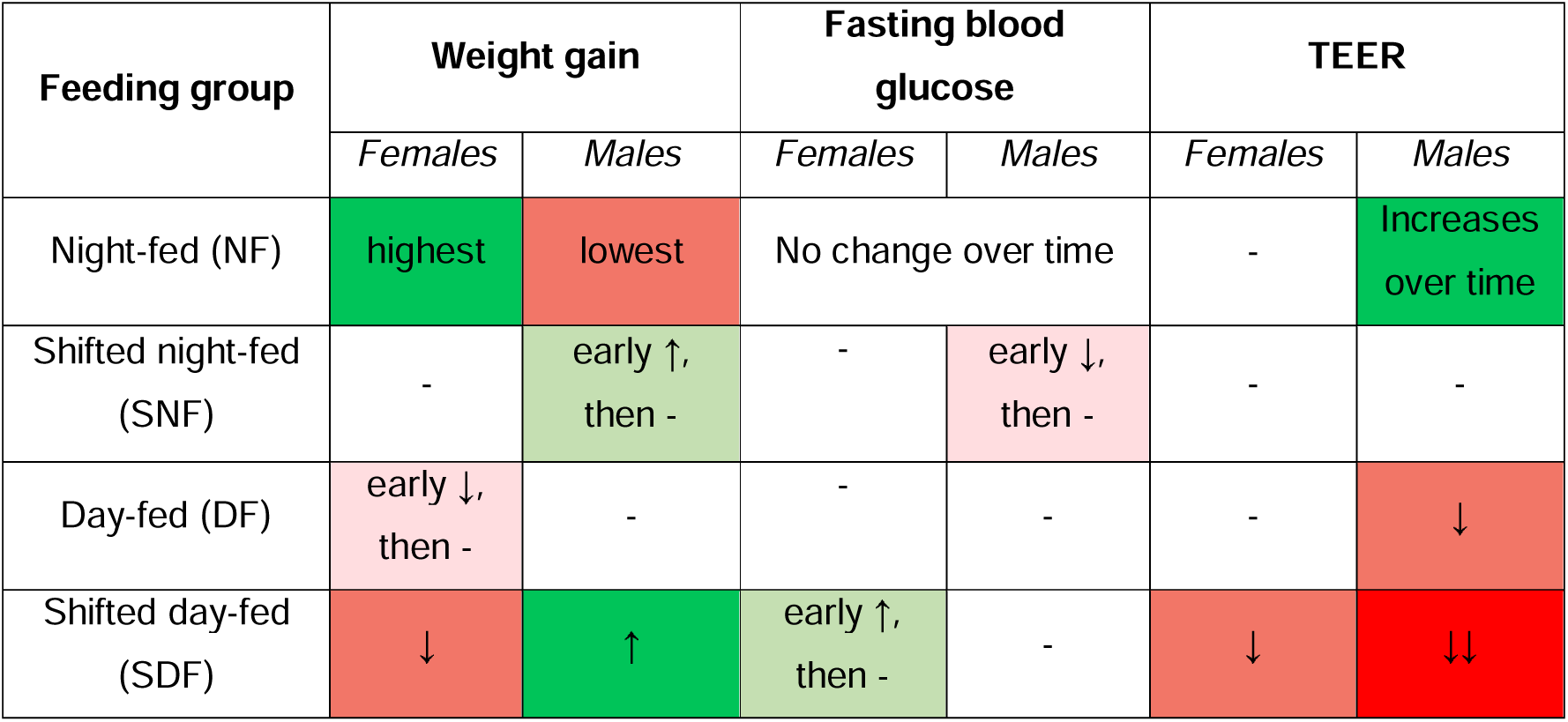
Cumulative changes in physiological parameters relative to the NF group.

| Feeding group | Weight gain |  | Fasting blood glucose |  | TEER |  |
| --- | --- | --- | --- | --- | --- | --- |
|  | *Females* | *Males* | *Females* | *Males* | *Females* | *Males* |
| Night-fed (NF) | highest | lowest | No change over time |  | - | Increases over time |
| Shifted night-fed (SNF) | - | early ↑, then - | - | early ↓, then - | - | - |
| Day-fed (DF) | early ↓, then - | - | - | - | - | ↓ |
| Shifted day-fed (SDF) | ↓ | ↑ | early ↑, then - | - | ↓ | ↓↓ |

Our data reveal the presence of sex-specific responses to TRF, especially regarding weight gain. When mice were fed during the lights-on phase (DF, SDF), weight gain slowed in females, but increased in males. Delay shifting the mealtime by 4 h, independently of the light-dark phase, led to a further weight gain increase in males (Fig 2A). Struggle for DF and SDF females to retain weight was also highlighted by the 16 h fasting period, leading to the lowest weights at the food anticipation timepoint (Fig 2B). Weight gain in DF and daytime-snacking male mice has been observed before [40, 78, 79], aligning with our observations; however, we are unaware of studies looking at weight gain in female mice under TRF. It has been observed that, under light phase feeding, the majority of physical activity still occurs during the dark phase [80–82], therefore, further measurements of locomotion and, ideally, firing rates within the central clock neurons would be needed to determine if DF and SDF females stay more active during the dark phase when no food is available to replenish the lost calories. TRF-induced sex-specific changes in food intake could also explain the observed differences, measuring individual consumption would be necessary to properly interpret the weight gain data reported here.

Fasting blood glucose levels also showed sex-specific adjustment to TRF. DF in female mice and SNF in male mice led to a significant decrease in fasting blood glucose levels, although these differences disappeared under longer entrainment (Fig 2C). This data contradicts previous observations of DF inducing higher fasting blood glucose [83], and suggests that glucose levels can normalize over long-term food entrainment. However, result interpretation would greatly benefit from measuring individual food intake and glucose levels at other timepoints.

Not much is known about TRF-driven changes in intestinal barrier function and epithelial resistance. Studies in this area often use disease models and have revealed that dark-phase TRF preserves intestinal integrity in mouse colitis model [84]. Our NF male mice also showed a significant increase in TEER over time (Fig 2D), complementing these observations. In contrast, circadian misalignment, caused by changes in the LD cycle and ‘wrong-time eating’, has been shown to downregulate the expression of tight junction proteins, increasing intestinal permeability [85, 86] and contributing to inflammatory bowel disease (for a review, see [87]). We also observed a significant decrease in TEER under prolonged SDF of both sexes and DF of males when food- and light-driven regulation would be misaligned (Fig 2D). Further measurements of expression of tight junction proteins and their upstream effectors (such as AMPK [88]), changes in microbiota, and TEER during acute fasting-refeeding (although fasting alone seems to have no effect on ileal permeability [89]) would aid in explaining how TRF regulates intestinal barrier function.

### Hepatic, gastric, and duodenal gene expression in response to TRF

Overall, we observed three distinct gene expression responses to food intake time, summarized in Table 3. For a gene to be classified as non-responsive, it had to show only one significant timepoint- or shift-specific expression difference for the whole duration of the experiment. Interestingly, although it was possible for genes to simultaneously show both timepoint- and shift-specific changes, they seemed to occur independently from each other.

**Table 3.**
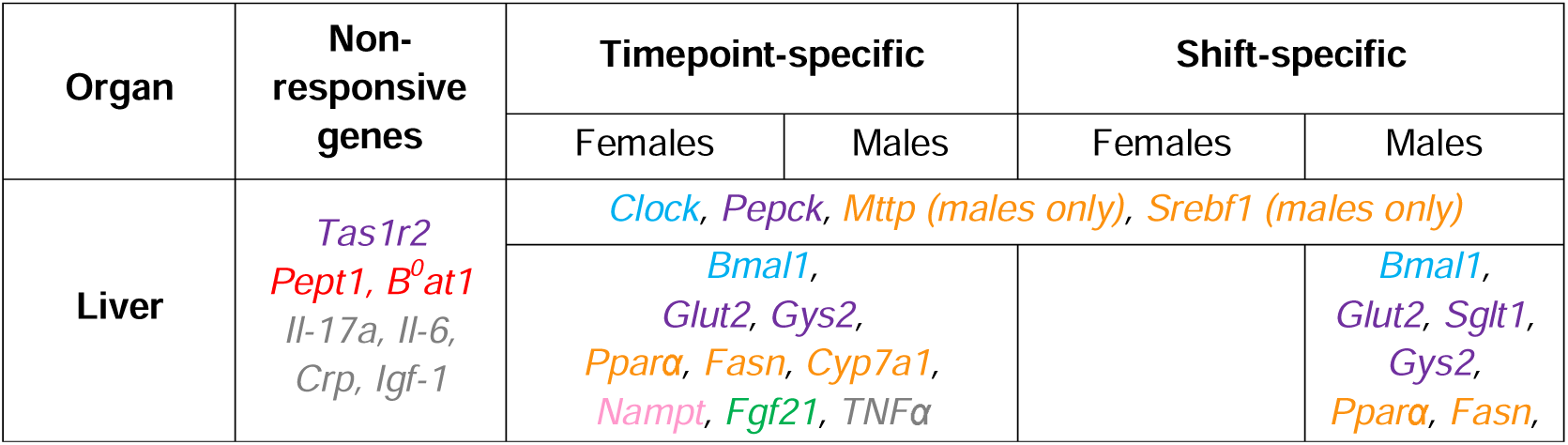
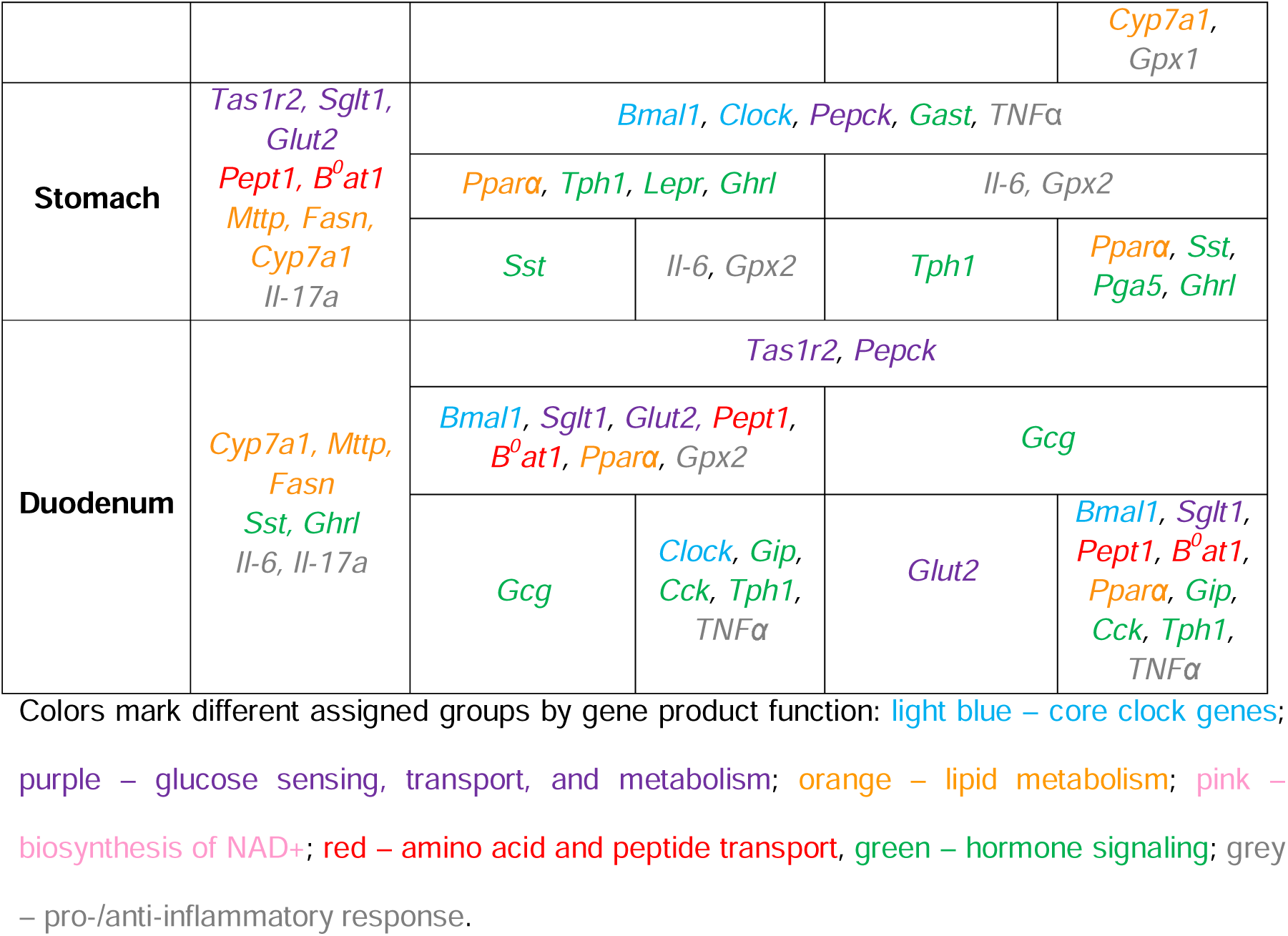
Summary of gene expression responses to food entrainment. Data pooled from all sacrifice days and TRF regimes.

Our data (Figs 3-6) led to three major conclusions: 1) digestive organs entrain to TRF at different rates; 2) DF drives an overall gene downregulation in the liver and an upregulation in the stomach and duodenum when compared to NF; 3) male and female mice can exhibit opposite timepoint-specific gene expression patterns, with males being more likely to also develop shift-specific responses.

Organ- and tissue-specific patterns in gene expression have been observed before, with surprisingly little overlap in cyclic genes and their phases, even if the tested genes are ubiquitously expressed and exposed to the same conditions [8, 22, 90–92]. This is similar to our observations, where the same gene could show timepoint-specific differences in one organ, but not in the others (e.g., *Pparα*) and where genes belonging to the same signaling pathway had no similarity in their responses (e.g., hepatic *Fgf21* and its upstream regulator *Pparα* [93]). At the same time, the gene *Pepck*, encoding a rate-limiting enzyme in gluconeogenesis, showed a similar timepoint-specific expression pattern in all organs, especially under SNF (Figs 4-6), demonstrating potential organ-to-organ communication. It appears that TRF is unable to synchronize timepoint-specific gene expression across the whole digestive system.

However, there was one major impact of TRF that spanned across two organs and all tested genes in both sexes. We did observe an overall gene upregulation in the stomach and duodenum and downregulation in the liver under DF and SDF (Figs 4-6). In the literature, such responses are rarely described, and their causes thus have not been discussed. A study by Deota et al. did show that TRF-specific gene up-/down-regulation patterns are similar between murine stomach and jejunum, but not liver and that specific gene groups can be upregulated in one organ, but downregulated in another [92], somewhat aligning with our observations. It has also been shown that fasting can induce gene expression changes in an organ-specific manner [94]; thus, further studies applying acute fasting and TRF in constant conditions would be needed to differentiate between fasting and TRF as causes for large-scale gene expression changes. In addition, transcriptomics and/or proteomics of multiple organs could provide further information on TRF-driven peripheral gene expression.

Although sex-specificity in circadian behavior has been reported before (for a review, see [95]), sex-specificity in TRF responses (or any other peripheral Zeitgeber, for that matter) has been understudied. Studies profiling metabolites have observed considerable sex-specific differences under TRF and other diets, especially in the liver [96, 97]. Also, the impact of fasting and refeeding on weight gain, energy expenditure, and regulation of appetite and nutrient metabolism has been shown to be sex-specific [98]. Our study revealed a considerable overlap of timepoint-specific responses in males and females; however, sex-specificity was strongly present in expression pattern changes over time and shift-specific responses. Long-term TRF studies with back- and-forth shifts would be beneficial to determine how fast males and females adjust to changes in food intake time and how stable the developed responses are. Such studies could also provide insights into the overall health impact of shifted or restricted feeding, which could be particularly important for shift workers and those often traveling across time zones.

Overall, our study demonstrates the complexity of TRF responses which are sex-, organ- and gene-specific and depend on the length of entrainment. It stresses the importance of including females in TRF and other Zeitgeber studies. Further details and integrative exploration of TRF under constant conditions and acute fasting-refeeding phases would be necessary to delineate the food-, fasting- and light-driven changes. In addition, increased sampling frequency would provide a more detailed timeline of the (sex-specific) adjustment to food intake time to fully unravel the interplay between sex, circadian rhythms, and food intake.

## Acknowledgments

We thank Single Cell and High Throughput Genomics Platform at Monash Health Translation project, especially Dr. Sen Wang, for performing the IFC analysis. We also thank Prof. Frédéric Gachon from the Institute of Bioscience at University of Queensland for the constructive exchange regarding experimental approaches and methodology.

## Competing interests

The authors of this study confirm there are no competing interests.

## Author contributions

All experiments were carried out at the Department of Anatomy and Physiology, The University of Melbourne, and the Florey Institute of Neuroscience and Mental Health. L.O-R., M.R.D., R.M., and J.B.F. designed the study and necessary experiments. M.R.D., L.O-R., R.M., B.H., A.K., M.R., T.E.F.C., and L.J.F carried out the experiments. L.O-R., M.R.D., R.M, and J.B.F. analyzed and interpreted the data. L.O-R., M.R.D., and R.M. wrote and critically revised the manuscript. All authors approved the final version of the manuscript. All persons qualifying for authorship are listed as such.

## Funding

This study was supported by the Swiss National Science Foundation Grant No. 187739 to L.O-R.

## Notes

### Competing Interest Statement

The authors have declared no competing interest.

### Summary of Updates

Result and discussion sections were updated, title change.

